# *In vitro* efficacy of Artemisia extracts against SARS-CoV-2

**DOI:** 10.1101/2021.02.14.431122

**Authors:** Chuanxiong Nie, Jakob Trimpert, Sooyeon Moon, Rainer Haag, Kerry Gilmore, Benedikt B. Kaufer, Peter H. Seeberger

## Abstract

Traditional medicines based on herbal extracts have been proposed as affordable treatments for patients suffering from coronavirus disease 2019 (COVID-19) caused by severe acute respiratory syndrome coronavirus 2 (SARS-CoV-2). Teas and drinks containing extracts of *Artemisia annua* and *Artemisia afra* have been widely used in Africa in efforts to prevent and fight COVID-19 infections. We sought to study the ability of different *A. annua* and *A. afra* extracts and the Covid-Organics drink produced in Madagascar to inhibit SARS-CoV-2 and feline coronavirus (FCoV) replication *in vitro.* Several extracts as well as Covid-Organics inhibit SARS-CoV-2 and FCoV replication at concentrations that did not affect cell viability. It remains unclear whether peak plasma concentrations in humans can reach levels needed to inhibit viral replication following consumption of teas or Covid-Organics. Clinical studies are required to evaluate the utility of these drinks for COVID-19 prevention or treatment in patients.

## 1. Introduction

Severe acute respiratory syndrome coronavirus 2 (SARS-CoV-2)[1] has caused a pandemic of coronavirus disease 2019 (COVID-19)[2–4] that resulted in a rising death toll as well as serious economic and societal consequences. Several vaccines were developed and approved in record speed, and are now being distributed as quickly as possible.[5,6] Still, affordable anti-viral treatments will still be needed for those that are not vaccinated or where vaccines fail to work. The clinical benefits of Remdesivir, the only antiviral drug approved for treatment of COVID-19, are being discussed controversially,[7] such that COVID-19 treatment remains largely supportive. Repurposing of drugs already licensed for other diseases is a comparatively fast method to meet the urgent need for effective antivirals against SARS-CoV-2. Artesunate **2** and other synthetic derivatives of the sesquiterpene lactone natural product artemisinin **1** that is isolated from *Artemisia annua* plants, are the key active pharmaceutical ingredient (API) of WHO-recommended anti-malaria combination therapies, used in millions of patients every year with few side effects.[8] Teas made from the leaves of *A. annua* plants are recommended in Traditional Chinese Medicine to treat malaria [9] and teas from *Artemisia* plants are widely used in many African countries to treat malaria patients, albeit contrary to WHO recommendations. [10]

Artemisinin and its synthetic derivatives have shown impressive effects against other parasitic infections,[11] a range of different types of cancer[12] and viruses [13] *in vitro* and in human clinical trials. In addition to pure, semi-synthetic substances, *Artemisia* teas and plant material have been explored for use to treat different diseases.[14] *A. annua* extract had shown anti-viral activity *in vitro* against SARS-CoV in 2005[15] and SARS-CoV-2 (EC50 value: 2.5 μg/mL)[16] [18]. Standardized *A. annua* extracts with high artemisinin content are currently being studied in an ongoing phase 2 clinical trial against COVID-19 [17]. Artesunate **2** was significantly more active than *A. annua* extracts against SARS-CoV-2 *in vitro* [16]. The excellent safety profiles of artemisinin-based drugs in humans and their ready availability for worldwide distribution at a relatively low cost render them attractive repurposing candidates for treatment of COVID-19.

Early during the pandemic, traditional medicines made from plant extracts were employed in various countries in efforts to prevent COVID-19 infections or treat COVID-19 patients. In South Africa, teas of *A. afra* that – in contrast to *A. annua* – do not contain artemisinin **1**, were used widely without *in vitro* or clinical data.[19] In Madagascar, Covid-Organics, a drink containing mainly *A. annua* extract and/or plant material, was announced by the President of Madagascar Andry Rajoelina as a “miracle cure” in April 2020, subsequently used in the country and exported to several African nations in hope to prevent and treat COVID-19 infections.[20] Due to fears that artemisinin combination therapies against malaria may become ineffective if artemisinin-based treatments are used against COVID-19, the WHO issued a warning against the use of traditional medicines.[21] More recently, the WHO modified its recommendation and called for an investigation into the potential efficacy of plant-based treatments.[22] To date, no *in vitro* data concerning anti-viral effects of *A. afra* plant extracts or Covid-Organics has been disclosed. Here, we report that various *A. annua* as well as *A. afra* plant extracts and Covid-Organics exhibit some anti-viral activity against SARS-CoV-2 *in vitro.* It is unclear whether the plasma concentrations that can be achieved in humans using such extracts are sufficient to prevent or treat COVID-19 infections. Human clinical trials will be required to answer the question whether the traditional medicines may indeed have an effect in preventing or treating COVID-19 infections.

## 2. Materials and Methods

### Plant Material

Dried leaves of *Artemisia* plants grown in different countries and in different years were obtained as donations (see supporting information). Covid-Organics was purchased in Madagascar.

### Extraction

Distilled water (10 mL) was heated to 90 °C in an Erlenmeyer flask. Dried plant material (1 g) was added to the solvent and kept for two minutes at 90 °C, then 20 minutes at room temperature. The mixture was filtered using filter paper and solid material washed with fresh distilled water (20 x 2 mL). The solvent was removed by rotary evaporation and the solid material was stored at −10 °C prior to sample preparation. An ethanolic extract of *A. annua var. CPQBA 1* was prepared by treating 100 g of dried leaves of *A. annua* with 1 L of ethanol at 70 °C for 20 days.

### Sample Preparation

Dried extract was warmed to room temperature before DMSO (3 mL) was added and the mixture heated (40 °C) to ensure solvation. The solution was filtered using a syringe filter and stored in a snap-close vial.

### Cell Culture

African green monkey kidney VeroE6 cells (ATCC CRL-1586) and Crandell-Rees Feline Kidney (CRFK, ATCC CCL-94) cells were maintained at 37 °C with 5% CO2 in Minimum Essential Medium (MEM; PAN Biotech, Aidenbach, Germany) supplemented with 10% fetal bovine serum (PAN Biotech), 100 IU/mL penicillin G and 100 μg/mL streptomycin (Carl Roth, Karlsruhe, Germany).

### Virus isolates

The SARS-CoV-2 BavPat 1 isolate (SARS-CoV-2/human/Germany/BavPat 1/2020) was provided by Dr. Daniela Niemeyer and Dr. Christian Drosten (Charité, Berlin, Germany) and obtained from an outbreak in Munich, Germany, in February 2020 (BetaCoV/Germany/BavPat1/2020). Feline coronavirus (ATCC VR-989, WSU 79-1683) was propagated and titrated on CRFK cells.[23]

### Plaque reduction antiviral assay

Tenfold dilutions of the compounds described above were prepared in cell culture medium. To determine the effect of the compounds, dilutions were incubated with 100 plaque forming units (PFU) of FCoV or SARS-CoV-2 for one hour at 37 °C. To determine the antiviral effect, tenfold dilutions of the compound-virus mix was incubated with CRFK or VeroE6 cells for 45 minutes at room temperature respectively. The cells were washed with PBS once, overlayed with 1.3% methylcellulose containing medium and plaque formation assessed two days post infection. Plaques of FCoV were stained with specific antibodies (primary antibody: mouse anti feline coronavirus monoclonal antibody at 1:500 dilution solution, Bio-Rad; secondary antibody: Alexa 488-labeled goat anti-mouse at a 1:500 dilution; Thermo Fisher) and counted manually by fluorescence microscopy (Axio observer, Zeiss). SARS-CoV-2 plaques were stained using crystal violet.

### Cell viability assays in VeroE6

To evaluate cytotoxic effects of the tested extracts, compounds and diluent (DMSO), cell viability was studied using Cell Counting Kit-8 (CCK8, Merck, Germany). VeroE6 cells were seeded at 10,000 cells per well of flat bottom 96-well plates (Thermo Fisher Scientific, Roskilde, Denmark). The next day, medium was exchanged containing specified concentrations of the samples. Each concentration or dilution was tested in three replicates; at least six nontreated control wells were included in the assay. After 24 h of incubation at 37 °C and 5% CO2, CCK8 Reagent (10 μL) was added per well and plates were incubated for 1 h at 37 °C, prior to recording absorbance at 450 nm using a FLUOstar OPTIMA 96-well plate reader (BMG LABTECH, Offenburg, Germany). Absorbance recorded in each well was related to the average absorbance of nontreated control wells to calculate the percentage of cell viability. Sigmoidal dose response curves were fitted and median cytotoxic concentration (CC50) values were calculated with GraphPad Prism 8.0.0.

## 3. Results

### 3.1. Extracts and compounds

*A. annua* and *A. afra* plants were extracted using distilled water at 90 °C for two minutes and then 20 minutes at room temperature. Solids were removed by filtration and the solvents evaporated. The extracted materials were dissolved in DMSO and filtered. Artemisinin (500 mg) was dissolved in DMSO (3 mL). Covid-Organics (50 mL) was dried by rotary evaporation and dissolved in DMSO (3 mL) (see supporting information for details).

### 3.2. Inhibition of feline coronavirus (FCoV) plaque formation by plant extracts and Covid-Organics

To assess the antiviral activity of the extracts and pure artemisinin, we first used the feline coronavirus (FCoV), a biological safety level (BSL)2 coronavirus related to SARS-CoV-2 that can be handled outside of a BSL3 facility. The extracts were diluted tenfold in DMEM medium and incubated with FCoV solution with 100 PFU for 45 min at room temperature. Virus and inhibitor mix were incubated with a monolayer of CRFK cells and plaque formation assessed two days post infection. A typical dose-dependent curve of virus inhibition is shown in Figure 2 (for details see the supporting information). All samples inhibited the virus at high concentrations. The antiviral activity was clearly reduced below 2 mg/mL, as no virus inhibition was observed, while all samples showed similar activity at the range of 5 – 10 mg/mL. The half maximal effective concentration (EC50) was estimated based on these curves (Table 2). Covid-Organics showed moderate inhibitory activity against FCoV (Figure S1, supporting information). The most promising samples based on the FCoV inhibition data were selected for in vitro inhibition studies using SARS-CoV-2 in the BSL3 facility.

**Table 1.** Selected dried leaves, extracts and Covid-Organics drink tested for activity. Year corresponds to when the plant material was harvested or extract/commercial drink was produced. For a full list of samples tested, please see supporting information.

| Type | Region | Year |
| --- | --- | --- |
| *A. annua* | USA | 2019 |
| *A. annua* | Chad | 2019 |
| *A. annua* | Burkina Faso | 2019 |
| *A. annua* var.<br>CPQBA 1<br>alcoholic<br>extract | Brazil | 2020 |
| *A. afra* | France | 2015 |
| Covid-Organics | Madagascar | 2020 |
| Artemisinin (**1**) | - | - |

**Table 2.** Inhibitory activity of the extracts against two different types of coronaviruses.

| Sample | EC<sub>50</sub>FCoV<br>(mg/mL) | EC<sub>50</sub>SARS-CoV-2<br>(mg/mL) | CC<sub>50</sub>VeroE6<br>(mg/mL) | Selectivity<br>index |
| --- | --- | --- | --- | --- |
| *A. annua* (Chad) | 7.65 ± 2.21 | 2.04 ± 0.61 | 14.05 ± 6.08 | 6.89 |
| *A. annua* (Burkina Faso) | 3.07 ± 0.70 | 0.88 ± 0.23 | 19.66 ± 11.00 | 22.34 |
| *A. afra* (France) | 4.10 ± 1.27 | 0.65 ± 0.21 | 17.48 ± 8.01 | 26.89 |
| *A. annu* (Brazil) | 12.99 ± 3.11 | 0.95 ± 0.33 | 19.54 ± 10.64 | 20.56 |
| Artemisinin | 6.69 ± 1.05 | 4.23 ± 1.88 | 18.18 ± 8.6 | 4.30 |
| *A. annua* (USA) | 13.02 ± 4.02 | 3.42 ± 0.66 | 12.04 ± 4.73 | 3.52 |

### 3.3. *In vitro* Inhibition of SARS-CoV-2 plaque formation by Artemisia extracts and Covid-Organics

The extracts with the highest activity in on FCoV were subsequently tested for their antiviral activity against SARS-CoV-2 using plaque reduction assays. SARS-CoV-2 plaque numbers were reduced following incubation with the extracts at at 1.83 mg/mL, while no activity was detected at 0.18 mg/mL. *A. annua* and *A. afra* aqueous extract as well as an *A. annua* ethanolic extract showed the strongest inhibition with EC50 values below 1 mg/mL (Figure 3, Table 2).

The cytotoxicity of all samples towards VeroE6 cells was assessed to ensure that any antiviral effects were not caused by toxicity. Using the Cell Counting Kit-8, the 50% cytotoxic concentration (CC50) of each extract in VeroE6 cells was determined and showed a similar toxicity profile with a CC50 around 10-20 mg/mL (Table 2). The selectivity index (SI) was assessed by comparing the EC50 against SARS-CoV-2 with the CC50 of the extracts. The aqueous extracts of *A. annua* from Burkina Faso and Brazil and *A. afra* from France showed an SI > 20. These tests confirmed that the extracts can inhibit the replication of SARS-CoV-2 at levels that are not toxic to the cells.

The Covid-Organics drink was first concentrated by rotary evaporation and then diluted for the test. As the exact mass concentration was unknown, the inhibitory activity is shown as percentage to the raw drinks (Table 3). The EC50 against SARS-CoV-2 was determined to be 7.73% of raw drink using the plaque reduction assay. The selectivity index was thereby 5.28.

**Table 3.**
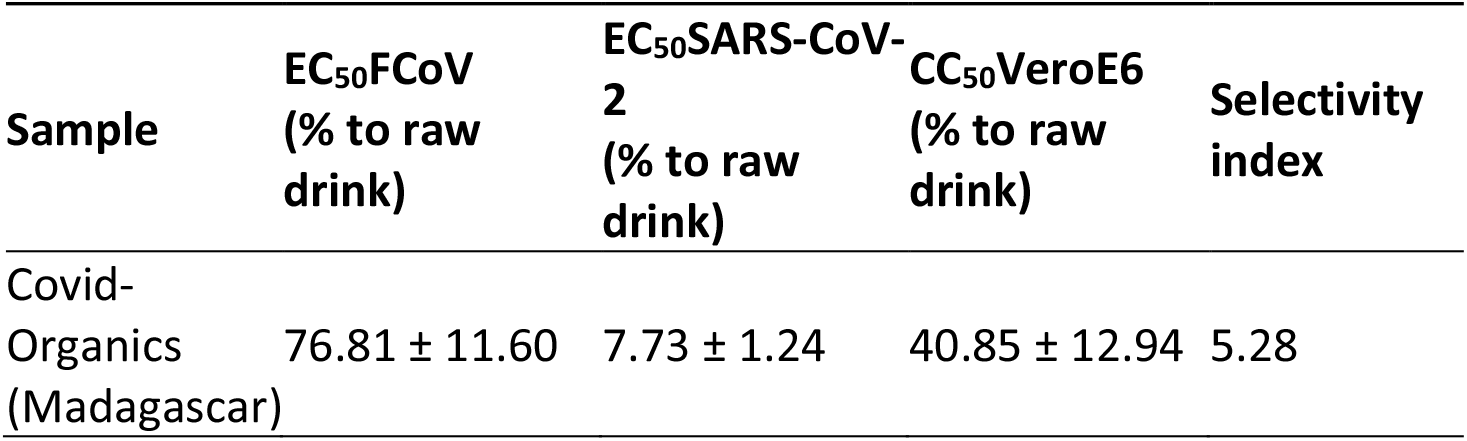
Inhibitory activity of Covid-Organics against two different coronaviruses.

| Sample | EC<sub>50</sub>FCoV<br>(% to raw drink) | EC<sub>50</sub>SARS-CoV-2<br>(% to raw drink) | CC<sub>50</sub>VeroE6<br>(% to raw drink) | Selectivity<br>index |
| --- | --- | --- | --- | --- |
| Covid-Organics (Madagascar) | 76.81 ± 11.60 | 7.73 ± 1.24 | 40.85 ± 12.94 | 5.28 |

## 4. Discussion

Natural products have been a rich source for the discovery of new drugs. For thousands of years humans have turned to plants in attempts to prevent, alleviate, or heal different diseases. Today, these “traditional medicines” have been broadly replaced by synthetic active pharmaceutical ingredients. Still, compounds obtained by isolation from plants are used in modern approaches to treat infectious diseases, cancer, and other ailments. Many natural products exhibit antiviral activities.[24] Artemisinin, isolated from *A. annua* plants, and the semi-synthetic derivative artesunate **2** obtained by chemical modification, inhibit a variety of viruses[13] including SARS-CoV.[15] Given the similarities between SARS-CoV and SARS-CoV-2, we investigated the use of artemisinin **1** (Figure 1), artemisinin derivatives such as artesunate **2**, as well as *A. annua* extracts for the inhibition of SARS-CoV-2 *in vitro* [16]. Artesunate was found to be most effective in inhibiting the virus, but *A. annua* extracts also revealed some activity *in vitro.* Human phase 2 clinical trials are currently ongoing to test the efficacy of these substances in treating COVID-19.[17] Other groups reported similar findings concerning pure synthetic artemisinin derivatives[25] and whole *A. annua* plant material.[18]

**Figure 1.**
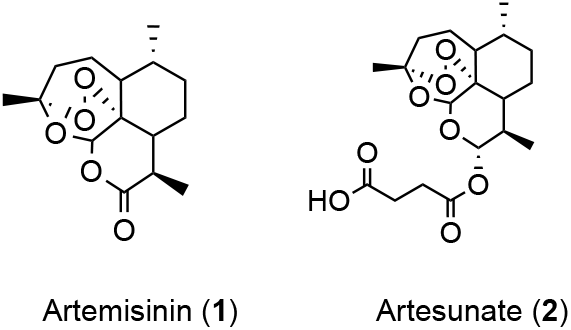
Chemical structures of artemisinin **1**, derived by isolation from *A. annua* plants and the semi-synthetic derivative artesunate **2**, that is the active pharmaceutical ingredient in WHO-recommended anti-malarial medications.

**Figure 2.**
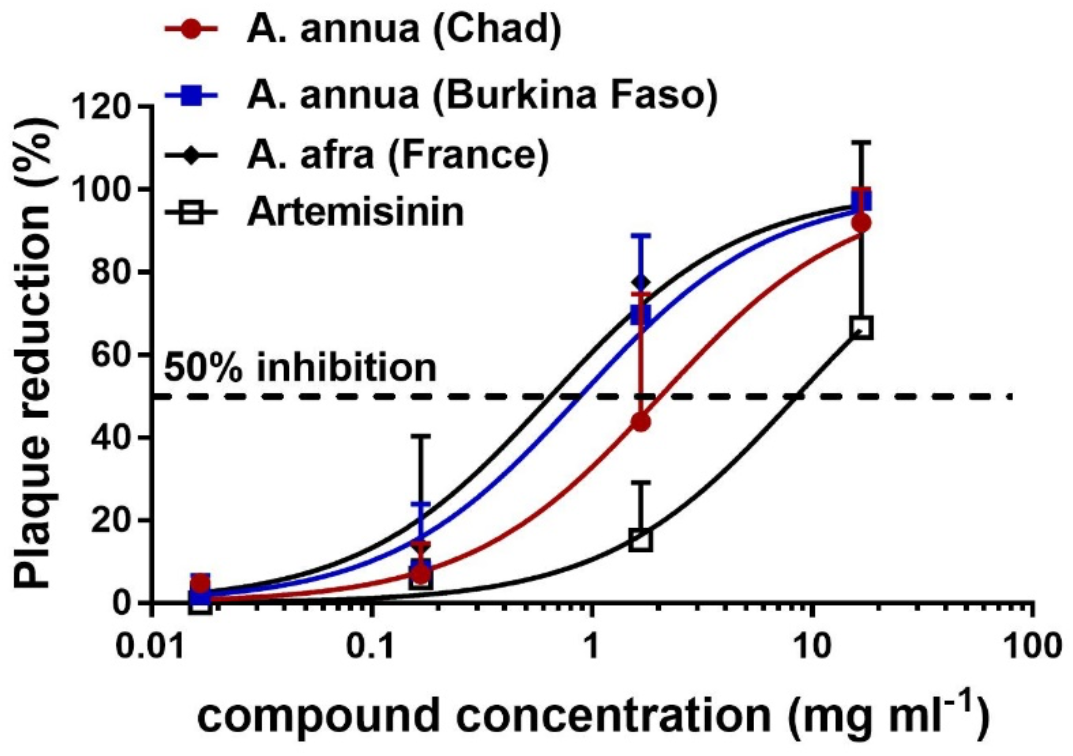
Dose-dependent inhibition of FCoV growth by addition of *Artemisia* extracts. Values are expressed as mean ±SD, n=3. For full dataset and experimental details, see supporting information.

**Figure 3.**
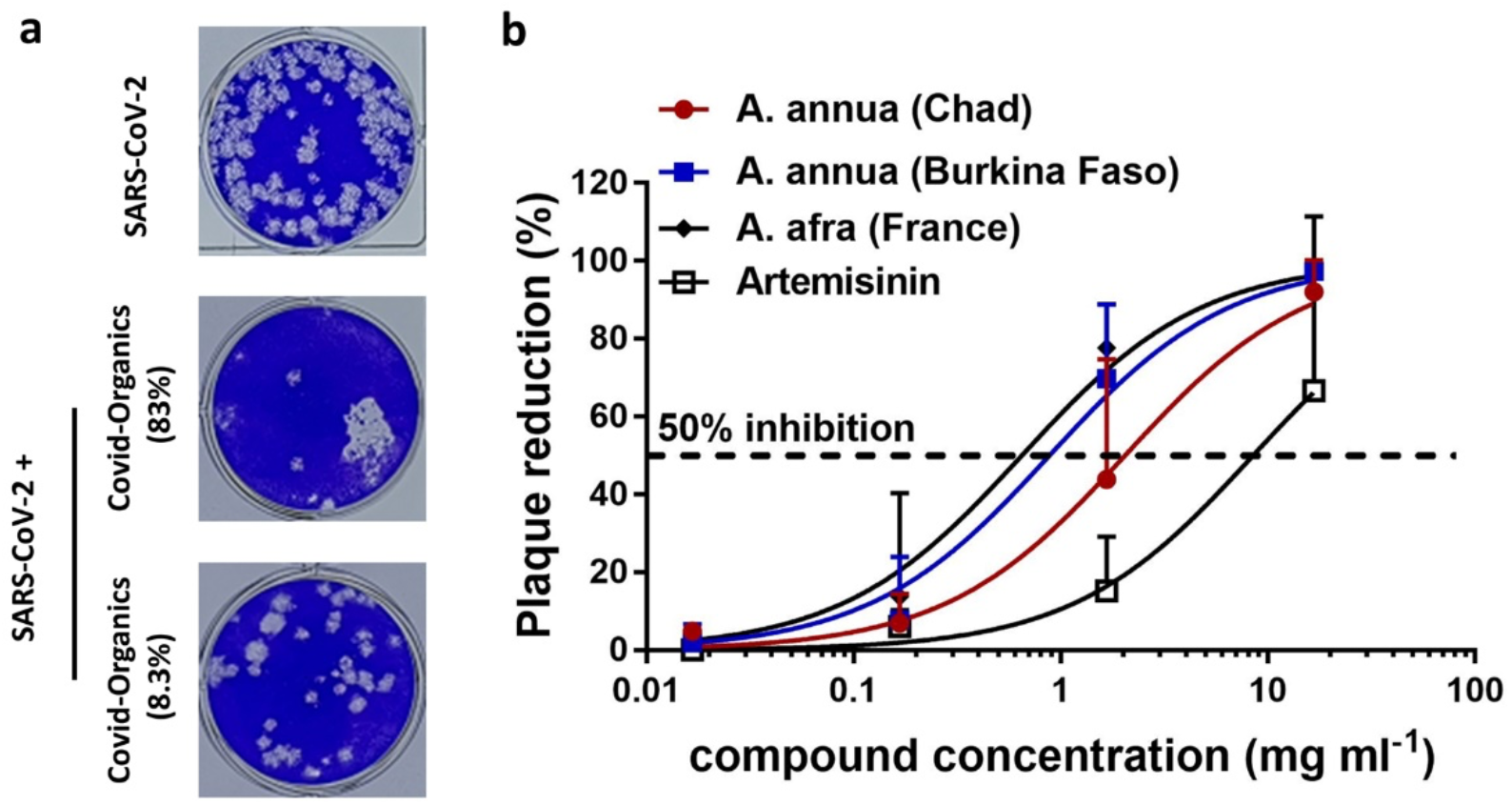
(a) Images of the SARS-CoV-2 plaques incubated with different dilutions of Covid-Organics. The dose is expressed by percentage of the raw drink. (b) Concentration-dependent inhibition SARS-CoV-2 replication using different extracts. Values are expressed as mean ±SD, n=3.

Reports that teas made from *A. annua* and *A. afra* plants,[19] as well as Covid-Organics, a drink made from *A. annua* leaves and *Ravensara aromatica* in Madagascar,[20] have been used throughout Africa. Our earlier observation that *A. annua* extracts can inhibit SARS-CoV-2 *in vitro* prompted us to test whether such extracts can indeed inhibit coronaviruses. To assess the inhibitory activity of the different plant extracts against two different coronaviruses, the feline FCoV and a human isolate of SARS-CoV-2, we performed plaque reduction assays *in vitro.* Screening of samples was performed using FCoV in a BSL-2 facility in blind fashion. The activity of the most promising samples was subsequently tested in a BSL3 facility using a SARS-CoV-2 isolate. Short pre-incubation with the extracts significantly inhibited plaque formation of both viruses, indicating that the extracts suppress viral replication in a dose dependent manner *in vitro* (Figure 3).

Some differences in the inhibitory activity against FCoV and SARS-CoV-2 were observed and confirm reports that SARS-CoV-2 is more sensitive to antivirals than FCoV. The extracts, with some slightly varying activities, were found to inhibit both coronavirus FCoV and SARS-CoV-2. Among the active samples were extracts of *A. annua* and *A. afra* plants (Figure 3). This finding was surprising considering that *A. afra* does not contain artemisinin and even more so since synthetic artesunate, a pharmacokinetically improved derivative of artemisinin, proved most active in previous *in vitro* SARS-CoV-2 inhibition studies [16, 25]. Our findings suggest that not just artemisinin but also other compounds present in the *Artemisia* extracts have inhibitory effects towards SARS-CoV-2. It has been suggested that flavonoids present in Artemisia species are active against SARS-CoV-2.[27]

The commercial Covid-Organics drink prepared from *A. annua* plants was promoted by the president of Madagascar in April 2020 at the beginning of the COVID-19 pandemic without any published scientific support as to its efficacy. Subsequently, a controversy around the product and its use in several countries in Africa arose due to fears that the use of artemisinin containing extracts may result in a resistance to this drug by *Plasmodium falciparum,* the parasite that causes malaria. We felt it was important to test the extracts *in vitro* and determine whether this product has any antiviral activity. In our *in vitro* test, Covid-Organics was found to inhibit SARS-CoV-2 at the concentrations we investigated (Figure 3), with an EC50 of 7.73% to raw drink and a selectivity index of 5.28. Covid-Organics exhibited a higher antiviral activity against SARS-CoV-2 compared to FCoV, which could be attributed the different interactions with the viruses and/or host cells. Even though inhibition of virus infection is noticed, the selectivity index is not very promising.

For all samples – *A. annua* and *A. afra* extracts as well as Covid-Organics – tested, it remains to be proven whether serum levels required to inhibit the virus can be reached in patients. Extracts showed some toxicity at higher concentrations but the selectivity index of 10 opens a useful therapeutic window to be explored in human clinical trials. It will have to be tested whether such extracts exhibit activity in the upper respiratory tract.

*Artemisia* extracts used as teas and Covid-Organics may be potential countermeasures for COVID-19. However, the *in vitro* inhibition data reported here needs to be followed up with investigations in animal models as well as human clinical trials before recommendations for use in patients can be made.

## Supplementary Materials

The following are available Figure S1: Inhibitory activity of Covid-Organics against two different coronaviruses. Values are expressed as mean ±SD, n=3. Table S1: List of dried leaves and compounds from various places and their inhibitory activity against FCoV.

## Author Contributions

Conceptualization, P.H.S., K.G. and B.B.K.; methodology, C.N. and J.T.; formal analysis, C.N. and J.T.; investigation, C.N. and J.T.; extraction, S.M.; sample preparation, S.M.; resources, P.H.S. and B.B.K.; writing—original draft preparation, review and editing, C.N., P.H.S. and B.B.K. with the input from all authors. All authors have read and agreed to the published version of the manuscript.

## Acknowledgments

We thank *ArtemiLife Inc.* for providing the *A. annua* plant material from Kentucky, Lucile Cornet-Vernet for *A. annua* and *A. afra* plant material, Prof. Pedro Melillo de Magalhaes (Univ. de Campinas, Brazil) for *A. annua* plant material and *A. annua* ethanolic extract, and Dr. H.M. Hirt for *A. annua* plant material. We thank an anonymous donor for providing an unopened bottle of Covid-Organics obtained in Madagascar. We thank Ann Reum for technical assistance.

