## Supporting Information for "*In vitro* efficacy of Artemisia extracts against SARS-CoV-2"

### SUPPLEMENTARY INFORMATION

#### 1. Reagents and Materials

Solvents were obtained from commercial suppliers and used without further purification. Information of dried leaves were listed below (Table S1). Samples are packed and stored under ambient conditions. For further details, please contact companies listed in the Table S1. Artemisinin was previously prepared and purified by crystallization using published protocols.[S1] Crystals were ground prior to use. Covid-Organics was used without further purification. Filter paper used was Rotilabo type 113A, diameter 240 mm, obtained from Carl Roth.

| Sample | Year | EC50 $_{FCoV}$ (mg/mL) | Affiliation |
|---|---|---|---|
| *A. annua* (Germany) | 2019 | 5.79 ± 1.48 | Teemana |
| *A. annua* (Nigeria) | 2019 | 8.38 ± 2.30 | Lucile Cornet Vernet |
| *A. annua* (Chad) | 2019 | 7.65 ± 2.21 | Lucile Cornet Vernet |
| *A. annua* (Madagascar) | 2019 | 7.15 ± 2.00 | Lucile Cornet Vernet |
| *A. tridentata* (USA, Utah) | 2019 | 3.39 ± 0.88 | Lucile Cornet Vernet |
| *A. absinthium* (France) | 2019 | 6.14 ± 1.73 | Lucile Cornet Vernet |
| *A. annua* (Togo) | 2019 | 5.11 ± 1.06 | Lucile Cornet Vernet |
| *A. annua* (Congo RDC) | 2020 | 2.94 ± 0.90 | Lucile Cornet Vernet |
| *A. annua* (France) | 2019 | 5.55 ± 1.26 | Lucile Cornet Vernet |
| *A. annua* (Burkina Faso) | 2019 | 3.07 ± 0.70 | Lucile Cornet Vernet |
| *A. afra* (Chad) | 2019 | 3.23 ± 0.49 | Lucile Cornet Vernet |
| A. afra (Benin) | 2019 | 2.45 ± 0.49 | Lucile Cornet Vernet |
| *A. afra* (France) | 2019 | 4.10 ± 1.27 | Lucile Cornet Vernet |
| *A. afra* (France) | 2015 | 4.64 ± 0.72 | Lucile Cornet Vernet |
| *A. annua* var. (Brazil) | 2020 | 12.99 ± 3.11 | University of Campinas |
| Artemisinin | - | 6.69 ± 1.05 | - |
| *A. annua* var. CPQBA 1 alcoholic extract (Brazil) | 2020 | 0.009 ± 0.002 | University of Campinas |
| *A. Annua* (USA, Kentucky) | 2019 | 13.02 ± 4.02 | ArtemiLife Inc. |
| Covid-Organics (Madagascar) | 2020 | 76.81 ± 11.60 (% to raw drink) | Malagasy Institute for Applied Research |

**Table S1.** List of dried leaves and compounds from various places and their inhibitory activity against FCoV.

#### 2. Extraction of *Artemisia* leaves

##### General procedure: Extraction using distilled water

Distilled water (10 mL, VWR) was added to an Erlenmeyer flask (50 mL) and heated to 90 ˚C using a hot plate. Dried leaf material (1 g) was added to the temperature-stable water and allowed to keep for two minutes at 90 ˚C and then 20 minutes at room temperature. The samples were filtered using filter paper and the solid material was washed with room temperature water (20 X 2 mL). The filtered solution was then dried using a rotary evaporator at least two hours. Dried samples were dissolved in DMSO (3 mL). Sonication needed for maximal solvation. The sample was filtered through a syringe filter (Chromafil® xtra RC0.45). The sample was stored at -10 °C until use.

For the alcoholic extract, tincture mother was prepared with the same material CPQBA 1, into a rate of 1:10: 100g of dried leaves of *A. annua* var. CPQBA 1 to 1000 mL of ethanol 70° shaking it once a day by 20 days. The alcoholic extract (2mL) was dried by rotary evaporation and the remaining substance was dissolved in DMSO (3 mL). The sample was stored at -10 °C until use.

**Artemisinin.** Artemisinin (500 mg) was dissolved in DMSO (3.0 mL). The solution was transferred with a 1000 µL pipette to a 1.5 mL Eppendorf snap-close vial. Concentration of example sample: 167 mg/mL. The sample was stored at -10 °C until use.

**Covid-Organics.** Covid-Organics (50 mL) was dried by rotary evaporation dissolved in DMSO (3 mL). The sample was stored at -10 °C until use.

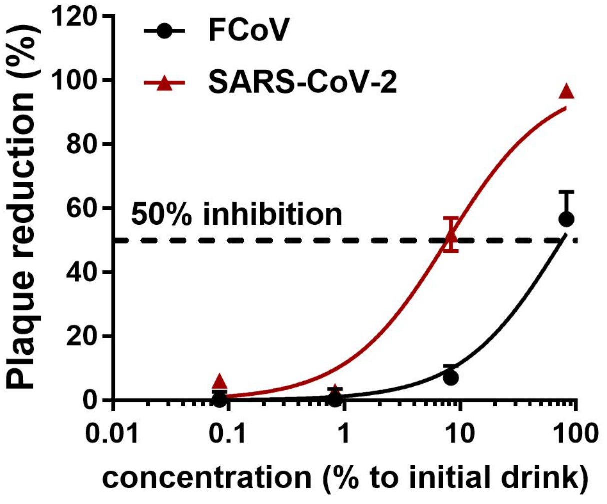

**Figure S1.** Inhibitory activity of Covid-Organics against two different coronaviruses. Values are expressed as mean ±SD, n=3.
